# Pre-clinical models of idiopathic scoliosis implicate sex-specific roles for complement activity in modulating spinal curve severity

**DOI:** 10.64898/2026.02.20.707049

**Authors:** Vida Erfani, Brian Ciruna

## Abstract

Idiopathic scoliosis (IS) is the most common spinal deformity and disproportionately affects adolescent females with severe disease. Although its etiology remains unclear, increasing evidence implicates inflammatory pathways in spinal curve progression. The complement cascade, a central component of innate immunity, has been linked to IS, but its functional role has not been directly tested. Here, we use zebrafish IS models to investigate the contribution of complement signaling to scoliosis pathogenesis. We develop novel genetic tools to upregulate or downregulate the core complement components *c3* and *c5*, enabling direct assessment of their roles in modulating spinal curvature. Applying these tools to the *sspo* IS model, we show that exogenous upregulation of *c3* significantly increases scoliosis severity in females. In *ptk7a* IS models, we demonstrate that loss of *c5* markedly exacerbates spinal curve severity specifically in males. Together, our findings identify complement signaling as a sex-dependent modifier of scoliosis severity in zebrafish and provide functional evidence linking innate immune activity to the progression of spinal deformity. These results offer a potential mechanistic explanation for the sex bias observed in human IS and highlight the complement cascade as a candidate target for therapeutic intervention.

## INTRODUCTION

Idiopathic scoliosis (IS) is a complex three-dimensional deformity of the spine defined by a lateral curvature of ≥10° (Cobb angle) accompanied by vertebral rotation, arising after birth in the absence of an identifiable underlying cause. Approximately 80% of cases present between 10 and 18 years of age (Cheng et al., 2015; Konieczny et al., 2013). When curvature develops during periods of rapid adolescent growth, progression can occur, with roughly 10% of cases advancing to a severity requiring surgical intervention (Miller, 1999). IS affects 1–4% of adolescents worldwide and disproportionately impacts females, who exhibit both higher prevalence and greater disease severity (Cheng et al., 2015; Konieczny et al., 2013).

The female bias in IS is robust and consistently reported yet remains poorly explained. During adolescence, the female-to-male ratio is approximately 3:1, increasing to as high as 10:1 in cases of severe curvature (Konieczny et al., 2013). Proposed explanations include sex-specific differences in spinal biomechanics and sexually dimorphic genetic susceptibility loci (Cheng et al., 2015). However, these hypotheses account for only a subset of cases and fail to provide a unifying mechanistic explanation. The biological bases of this pronounced sexual dimorphism remain to be elucidated.

Historically, IS has primarily been treated as a structural disorder, limiting therapeutic options to mechanical interventions such as bracing and spinal fusion (Hasler, 2013). These approaches can be socially burdensome, invasive, and associated with chronic morbidity. Progress toward less invasive and more targeted therapies requires a deeper understanding of the biological mechanisms underlying disease onset and progression. Genome-wide association and familial studies indicate that IS is a multifactorial disorder influenced by diverse genetic and environmental factors (Kou et al., 2019). No single genetic variant has emerged as a sole driver of disease, underscoring the complexity and incomplete nature of current etiological models.

Animal models like the zebrafish have emerged as powerful systems for studying axial deformities. There are numerous zebrafish models of IS that are mutant for biologically diverse gene functions yet converge on similar scoliotic phenotypes. (Bagnat and Gray, 2020; Boswell and Ciruna, 2017). Comparative analyses reveal model-specific defects but also shared pathophysiological features, suggesting the presence of common downstream mechanisms. One such shared feature is increased neuroinflammation, which has been reported in *sspo* (Rose et al., 2020), *ptk7a* (Van Gennip et al., 2018), and *rpgrip1l* (Djebar et al., 2024) mutant models.

Mutations in *ptk7a* and *rpgrip1l* disrupt ciliogenesis, which impairs motile-cilia mediated cerebrospinal fluid (CSF) flow (Djebar et al., 2024; Grimes et al., 2016; Van Gennip et al., 2018) and leads to defects in the assembly and maintenance of Reissner’s fibre—a glycoprotein filament extending through the ventricular cavities of the brain and spinal cord that is essential for CSF homeostasis (Rose et al., 2020). *Sspo* mutants also exhibit bulk CSF flow defects and loss of Reissner fibre formation in the absence of ciliary dysfunction, further implicating disrupted CSF homeostasis in IS pathogenesis (Rose et al., 2020; Troutwine et al., 2020). Notably, clinical imaging studies of IS patients corroborate a potential role for altered CSF dynamics in scoliosis (Algin et al., 2022; Tomita et al., 2024). Importantly, zebrafish studies indicate that CSF homeostasis defects ultimately cause neuroinflammation, and that inflammatory signals are both necessary and sufficient for the induction of scoliosis (Djebar et al., 2024; Rose et al., 2020; Van Gennip et al., 2018).

Across these zebrafish IS models, components of the complement system consistently rank among the most highly upregulated transcripts and proteins. Complement proteins serve diverse immunological, regenerative, and developmental functions; however, excessive or chronic activation promotes inflammation and tissue damage (Mastellos and Lambris, 2002; Merle et al., 2015). This raises a critical unresolved question: does complement activation contribute to the onset or severe progression of idiopathic scoliosis?

Complement activation is a highly conserved vertebrate immune mechanism (Zarkadis et al., 2001). It proceeds through a proteolytic cascade involving core components C1–C9, amplifying innate and adaptive immune responses, promoting inflammation, and culminating in formation of the membrane attack complex (MAC) (Merle et al., 2015). C3 occupies a central role in this cascade, mediating opsonization and generating pro-inflammatory anaphylatoxins. Its activity is tightly regulated by complement control proteins, including zebrafish regulators such as Rca2.1 and Rca2.2. Downstream, C5 initiates MAC formation (Podack et al., 1980) and also functions as a potent anaphylatoxin (Morgan, 2016). Dysregulation at either node could plausibly drive pathological inflammation within the spinal axis.

Although neuroinflammation has not yet been directly characterized in IS patients, multiple inflammatory markers correlate with disease incidence and severity (Bertelè et al., 2024; Makino et al., 2019; Shen et al., 2019; Wang et al., 2021). Complement-related proteins—including C3, C5, and the regulatory protein Factor H—are among the most upregulated serum factors reported in adolescents with IS (Lyu et al., 2026; Makino et al., 2019; Wang et al., 2021). Additionally, abnormal deposition of the complement inhibitor CD59 has been observed in intervertebral discs of IS patients (Teixeira et al., 2021). Collectively, these findings implicate complement dysregulation in IS but do not resolve whether it contributes to pathogenesis or arises secondary to other instigating factors.

Here, we directly test the necessity and sufficiency of the central complement components C3 and C5 in scoliosis induction and progression using zebrafish models of IS. By functionally manipulating complement activity in vivo, we aim to determine whether innate immune signaling acts as a modifier of spinal curvature and whether it contributes to the sex bias observed in disease severity.

## RESULTS

### Distinguishing abnormal from normal spinal curvatures in zebrafish

Zebrafish spines normally display a slight kyphosis within the precaudal region, analogous to the natural anatomy of the human spine. Abnormal curvatures are classified as departures from this norm. However, because the penetrance and expressivity of spinal curvatures in zebrafish IS models is variable (Djebar et al., 2024; Hayes et al., 2014; Meyer-Miner et al., 2022; Pumputis et al., 2025; Rose et al., 2020), diagnosing scoliosis penetrance can prove difficult when the phenotype is very mild.

In humans, the threshold for scoliosis diagnosis is a Cobb angle of 10 degrees or more. To empirically establish a diagnostic threshold for scoliosis in zebrafish, we performed micro-CT imaging on wildtype adult fish and quantified the angles of normal spinal curvature in both dorsovental (DV) and mediolateral (ML) planes. We found that the natural curvature of the precaudal spine in wildtype fish is more pronounced in females than males, with spinal curvatures in the DV plane ranging up to 24° in males and 31° in females. (Fig. 1). Therefore, the diagnostic thresholds for scoliosis should be sex-dependent. Notably, zebrafish spines do not normally curve in the ML plane, therefore these measurements represented total Cobb angle values for each fish. For the purposes of this study, these total Cobb angle values (male=24°, female=31°) were used as sex-specific thresholds for diagnosing scoliosis (Fig. S1).

**Figure 1:**
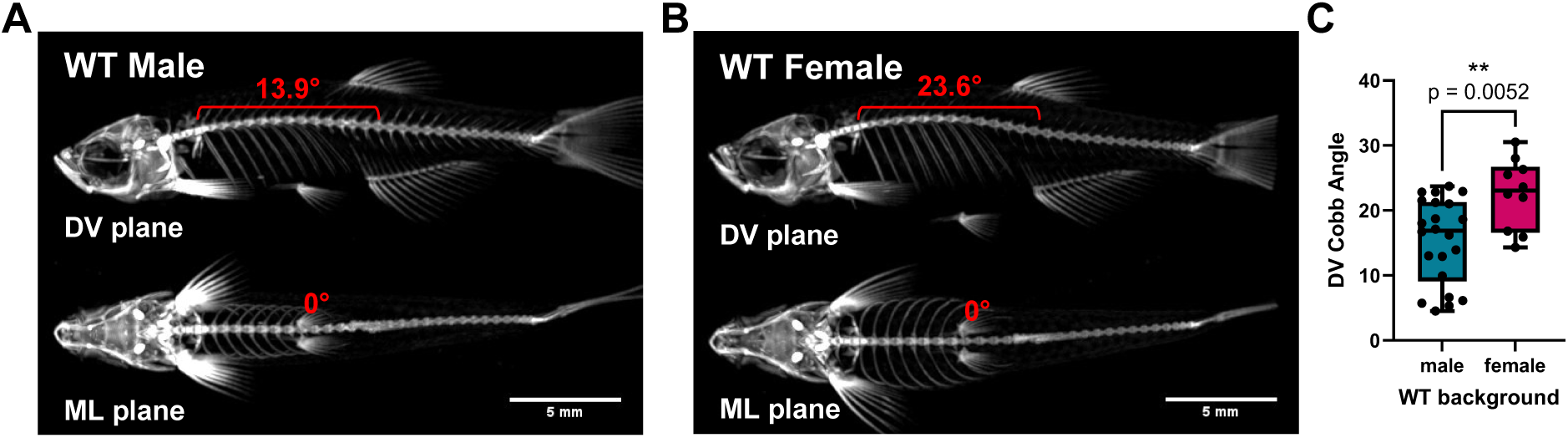
In wildtype zebrafish, females exhibit more pronounced natural precaudal spinal curvatures. Wildtype zebrafish display 1 spinal curvature within the DV plane, and 0 curvatures within the ML plane. Cobb angle measurements performed on micro-CT images of wildtype male (A) and female (B) adult zebrafish (5mpf) reveals a significantly greater degree of curvature in female fish than male siblings (C). *female*: n=10, mean: 22.5°, median=23.1, SD=5.4, max value: 30.5; *male*: n=22, mean:15.4, median:16.9, SD:6.5, max value: 23.7; p-value=0.0052, by unpaired two-tailed t test.

### Neuronal overexpression of *c3* does not cause scoliosis, but modestly increases spinal curve severity in female *sspo(dmh4/+) IS* models

The zebrafish genome contains 8 *c3* paralogues resulting from gene duplication events and speciation (Forn-Cuní et al., 2014; Gongora et al., 1998). It is speculated that many of these encoded proteins functionally overlap, based on high sequence complementarity and protein domain conservation (Forn-Cuní et al., 2014; Najafpour et al., 2020). In addition to their canonical roles, such as opsonization and chemotaxis, it has been suggested that zebrafish *c3 group b* genes (*c3b.1* and *c3b.2*) also play a unique role in the regulation and modulation of inflammation (Forn-Cuní et al., 2014). As outlined in Fig. S2, previous RNA-sequencing data demonstrated strong upregulation of *c3 group b* genes in the brains of severely scoliotic juvenile *sspo(dmh4*/*+)* fish (Rose et al., 2020). To investigate the functional consequence of elevated *c3* expression, we cloned and ectopically overexpressed *c3b.2* (here on referred to as *c3*) under a pan-neuronal promoter [*Tg(elavl3::c3*)] and screened for its effects on spine morphogenesis (Fig. 2A,B).

**Figure 2:**
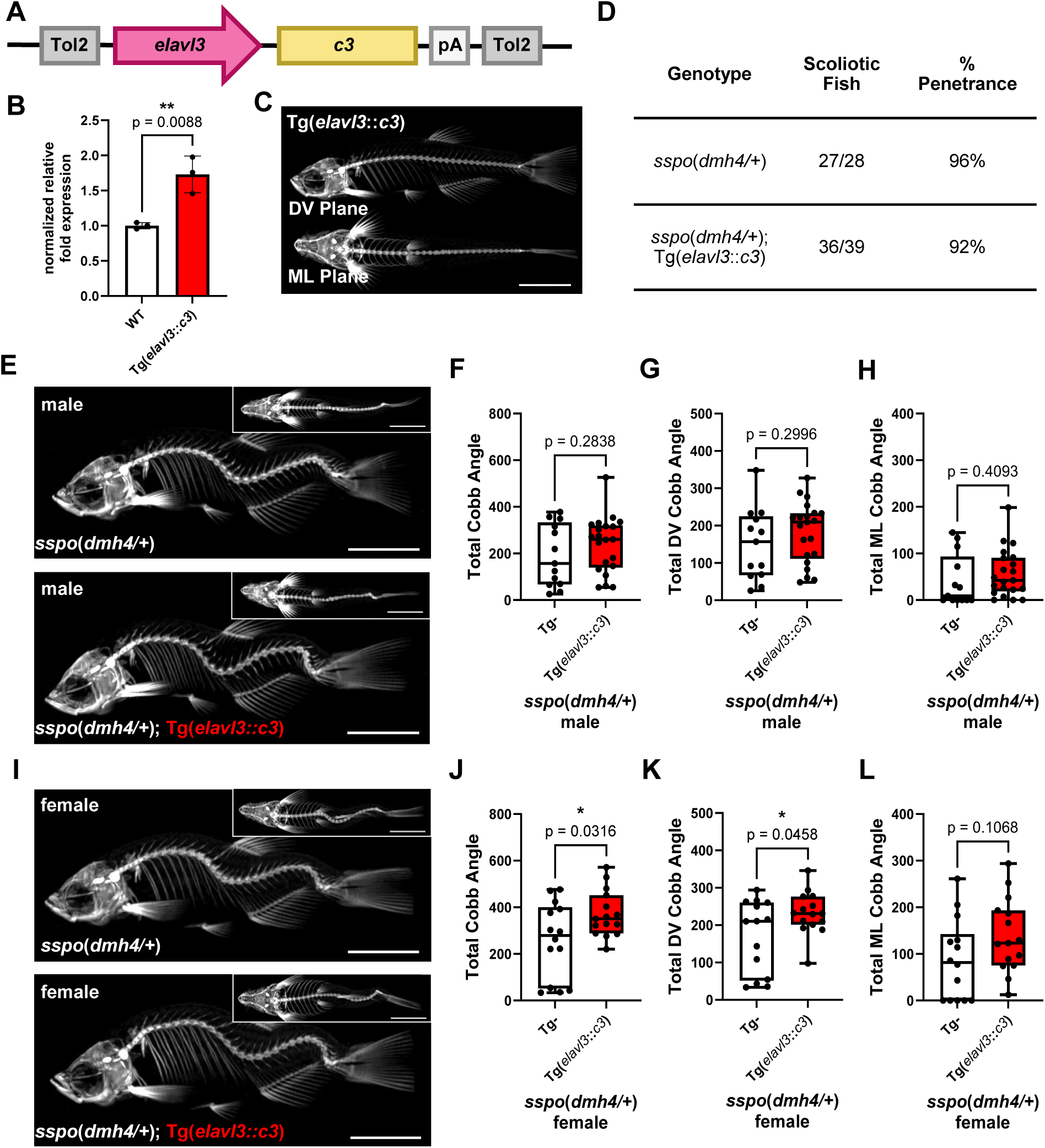
Pan-neuronal overexpression of *c3* in the *sspo*(*dmh4/+*) IS model leads to a female-specific increase in spinal curve severity. (A) A Tg(*elavl3::c3*) transgene driving pan-neuronal overexpression of *c3* was cloned, and (B) expression levels in stable transgenic animals validated by qRT-PCR. (C) Tg(*elavl3::c3*) expression does not cause axial phenotypes in a wildtype background. (D) Quantification of scoliosis penetrance in *sspo*(*dmh4/+*) and *sspo*(*dmh4/+*); Tg(*elavl3::c3*) siblings revealed no significant difference between cohorts (Two-sided Fisher’s exact test, p=0.6346; if separated by sex: females, p>0.9999; males, p>0.9999). To quantify the effect of c3 overexpression on *sspo*(*dmh4/+*) scoliosis severity in male (E) and female (I) fish, uCT imaging was performed, followed by Cobb angle analysis (F-H, J-L). The scoliosis phenotype was not penetrant in 4 fish (Tg-male, n=1; Tg+ male, n=2; Tg+ female, n=1). These data points have been eliminated from the analysis of scoliosis severity. Scale bars = 5mm.

*Tg(elavl3::c3*) expression was not sufficient to induce IS-like spinal curvatures in wildtype zebrafish, which fails to support a role for elevated complement activity in driving axial deformities (Fig. 2C). To assess for possible modifying effects of *c3* expression on existing scoliosis phenotypes, *Tg(elavl3::c3*) was crossed into the *sspo(dmh4*/+) genetic background. No significant difference in the penetrance of scoliosis was observed between *sspo(dmh4/+)* transgenic and non-transgenic siblings (Fig. 2D). However, a mild yet statistically significant increase in the severity of spinal curvatures was observed specifically in female *sspo(dmh4/+)*; *Tg(elavl3::c3*) fish, particularly in the ML plane (Fig. 2E-L). Consistent with published studies (Rose et al., 2020), a sex bias in scoliosis severity was not observed between non-transgenic *sspo(dmh4/+)* male and female control groups (Fig. S3). Therefore, the observation that female *sspo(dmh4*/+); Tg(*elavl3::c3*) fish exhibit a more severe total Cobb angle value relative to their male counterparts suggests a sex-dependent role for elevated C3 activity in modifying scoliosis phenotypes.

### Modulation of *rca2.1* expression levels does not significantly affect the *sspo(dmh4/+)* scoliosis phenotype

To further investigate the role for C3 in modulating scoliosis phenotypes, we attempted to increase endogenous C3 activity. Using CRISPR/Cas9 genome editing approaches, we generated stable transcriptless deletion alleles for *rca2.1* and the closely related zebrafish paralogue *rca2.2* (figure 3A,E) - negative regulators of complement activation (Tsujikura et al., 2015; Wu et al., 2012). We hypothesized that zebrafish *rca2.1* and *rca2.2* mutants would exhibit increased C3 protein deposits on autologous tissue, as has previously been shown in carp using antibody-mediated inhibition of RCA2.1/Tecrem function (Tsujikura et al., 2015). However, our studies of *rca2.1* and *rca2.2* loss-of-function phenotypes were constrained by our finding that homozygous deletion of *rca2.1* and homozygous co-deletion of *rca2.1* and *rca2.2* (*Df(Chr23:rca2.2, rca2.1)*) both resulted in larval lethality by 13 days post fertilization, prior to the onset of scoliosis in zebrafish IS models (Fig. 3C). Smaller indel mutations in *rca2.1* were generated but were also homozygous lethal (Fig. S4). Our studies were therefore limited to assessing scoliosis in *sspo(dmh4/+); rca2.1(del/+)* and *sspo(dmh4/+); Df(Chr23:rca2.2, rca2.1)/+* fish compared to *sspo(dmh4*/+) control siblings. Although we did not observe any significant change in scoliosis penetrance or scoliosis severity in these mutants (Fig. 3D,F), we cannot draw strong conclusions as *rca2.1* and *rca2.2* may be haplosufficient.

**Figure 3:**
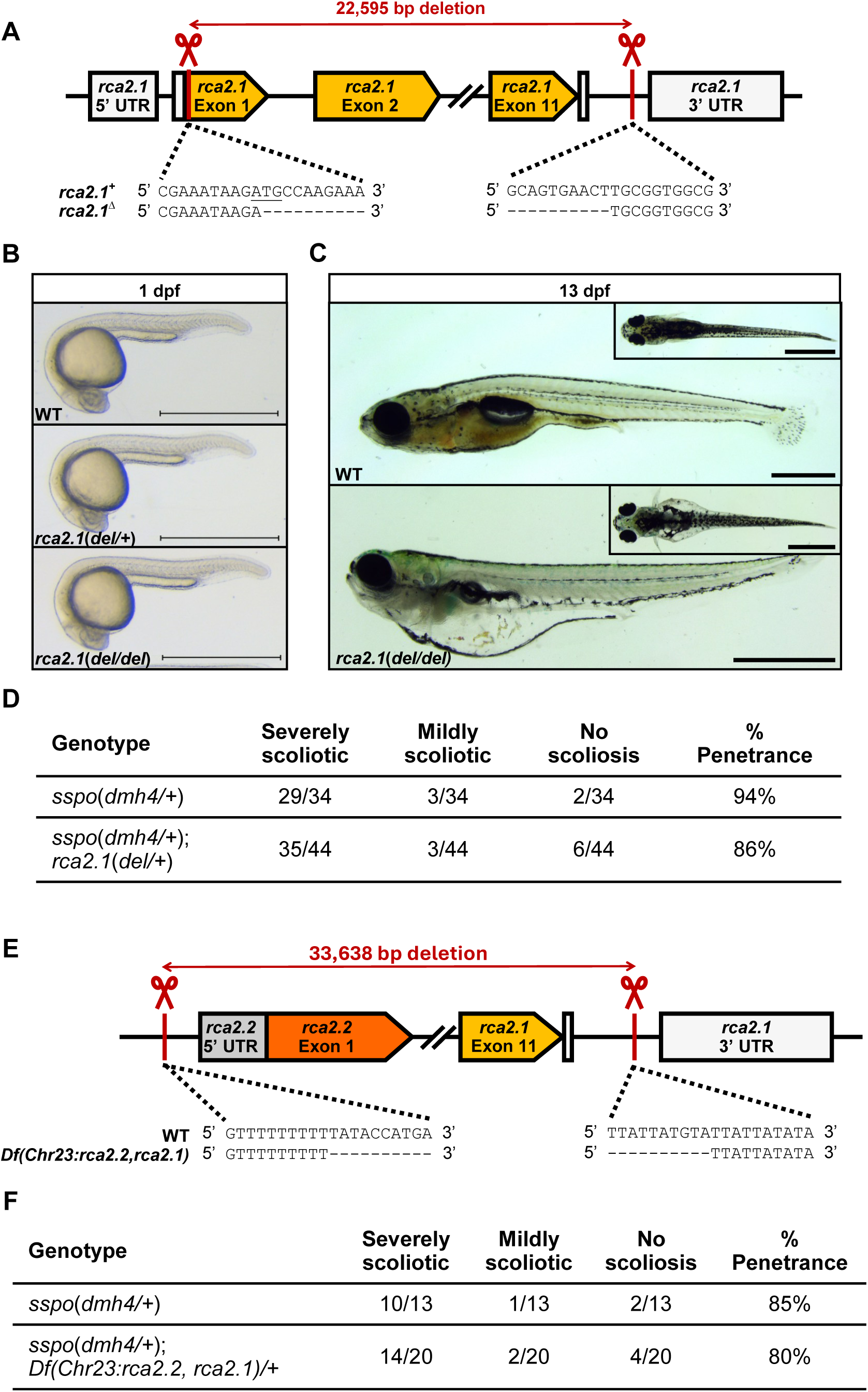
Heterozygous deletion of *rca2.1* and *rca2.2* does not cause scoliotic curvatures or affect scoliosis incidence in the *sspo(dmh4/+)* IS model. (A) A large deletion of the *rca2.1* open reading frame was generated using CRISPR/Cas9 genome editing. If a mutant transcript is formed, the predicted protein product would be: MRWRLKT*. (B-C) Heterozygote *rca2.1(del/+)* animals do not exhibit obvious developmental phenotypes while homozygous *rca2.1(del/del)* mutants experience severe pericardial edemas, growth retardation, and lethality by 2 wpf; Scale bars = 1mm. (D) Scoliosis penetrance and severity were qualitatively scored, revealing no significant difference in the proportion of phenotypic outcomes between *sspo(dmh4/+)* and *rca2.1(del/+)*; *sspo(dmh4*/+) siblings (Fisher’s exact test, p=0.6688). (E) A large deletion encompassing the syntenic *rca2.1* and *rca2.2* genes was generated. (F) Scoliosis penetrance and severity were qualitatively scored, revealing no significant difference in the proportion of phenotypic outcomes between *sspo(dmh4/+)* and *sspo(dmh4/+); Df(Chr23:rca2.2, rca2.1)/+* siblings (Fisher’s exact test, p>0.9999).

Conversely, we took an indirect approach to globally reduce activation of endogenous C3 proteins. To do this, we cloned and overexpressed the zebrafish negative regulator of complement activation *rca2.1* under a ubiquitous promoter [Tg(*Bactin2::rca2.1*)] (Fig. 4A,B). Stable Tg(*Bactin2::rca2.1*) fish did not display obvious axial defects (Fig. 4C).

**Figure 4:**
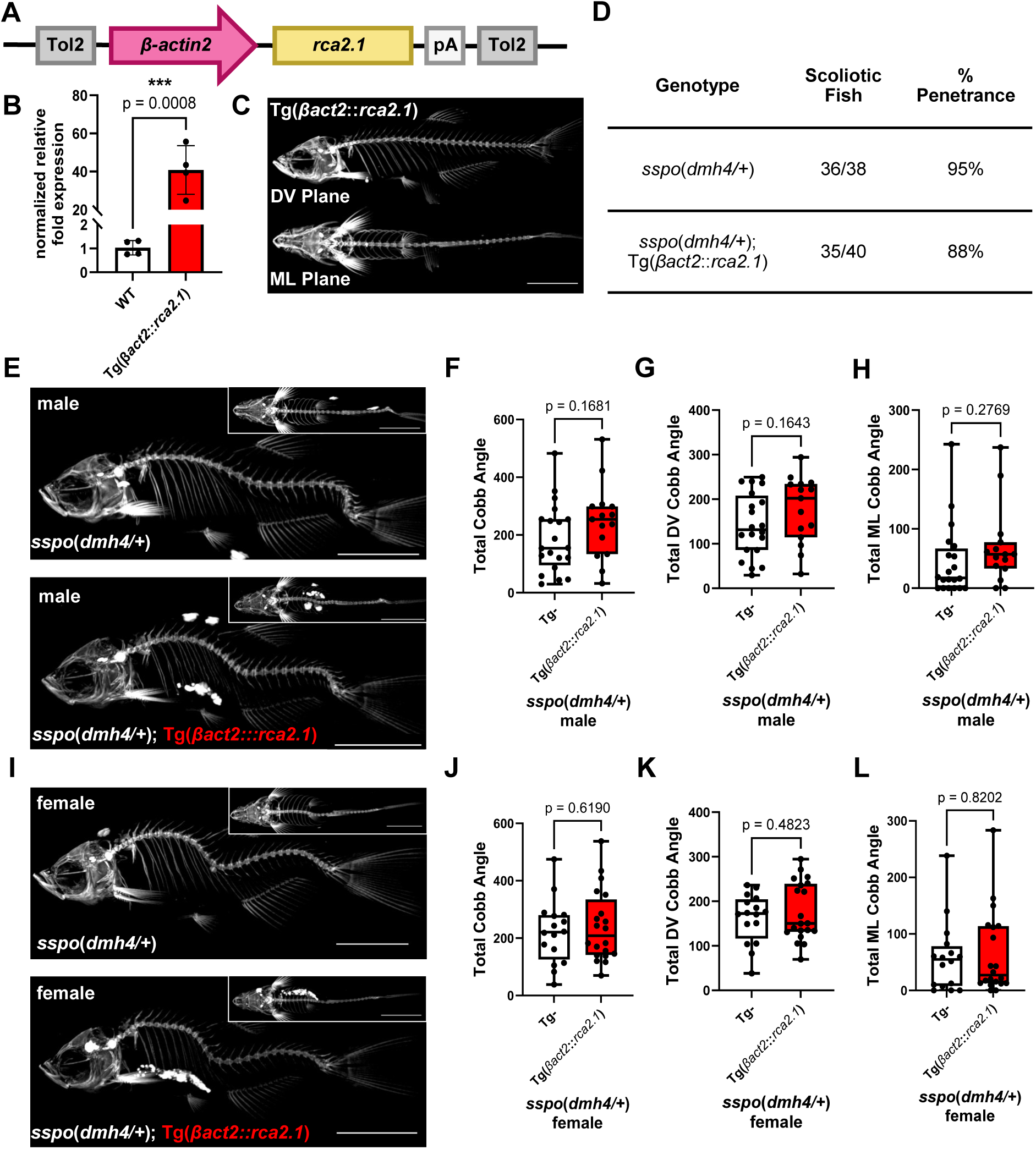
Overexpression of *rca2.1* in *sspo(dmh4/+)* IS models does not modulate spinal curve severity. (A) A Tg(*βactin2::rca2.1*) transgene driving ubiquitous overexpression of *rca2.1* was generated, and (B) expression levels in stable transgenic animals was validated by qRT-PCR. (C) Tg(*βactin2::rca2.1*) expression does not cause axial phenotypes in a wildtype background. (D) Quantification of scoliosis penetrance in *sspo*(*dmh4/+*) and *sspo*(*dmh4/+*);Tg(*βactin2::rca2.1*) siblings revealed no significant difference between cohorts (Two-sided Fisher’s exact test, p=0.4321; if separated by sex: females, p=0.6235; males, p=0.5768). To quantify the effect of *rca2.1* overexpression on *sspo(dmh4/+)* scoliosis severity, uCT imaging was performed on males (E) and females (I), followed by Cobb angle analysis (F-H, J-L). The scoliosis phenotype was not penetrant in 7 fish (Tg-male, n=1; Tg+ male, n=2; Tg-female, n=1; Tg+ female, n=3). These data points have been eliminated from the analysis of scoliosis severity. Scale bars = 5mm.

Additionally, we observed no significant difference in scoliosis penetrance between *sspo(dmh4/+);* Tg(*Bactin2::rca2.1*) fish and non-transgenic control siblings (Fig. 4D). Furthermore, Cobb angle analysis revealed no difference in scoliosis severity between these groups in either plane of rotation (Fig. 4E-L). We conclude that ubiquitous overexpression of *rca2.1* is not sufficient to impact scoliosis incidence or spinal curve severity in the *sspo(dmh4/+)* model.

### Deletion of *c5* does not influence the *sspo*(*dmh4/+*) scoliosis phenotype

As outlined in Fig. S2, RNA-sequencing results from a previous study revealed an upregulation of *c6* and *clu* in *sspo(dmh4/+)* fish (Rose et al., 2020) which are part of the terminal complement pathway, involved in MAC formation and regulation. To elucidate the role for the MAC in modulating zebrafish scoliosis phenotypes, we targeted *c5* which is necessary for initiation of MAC assembly (Podack et al., 1980). Using CRISPR/Cas9 genome editing, we generated a stable deletion allele of *c5* (Fig. 5A) and confirmed that mutants exhibited reduced cell lysis activity by hemolysis assay (Fig. 5B). Stable homozygous *c5* deletion did not result in any obvious axial defects (Fig. 5C). Scoliosis penetrance was scored, and we observed no significant difference between *sspo(dmh4/+)*; *c5(del/del)* fish and *sspo(dmh4/+)*; *c5(del/+)* control siblings (Fig. 5D).

**Figure 5:**
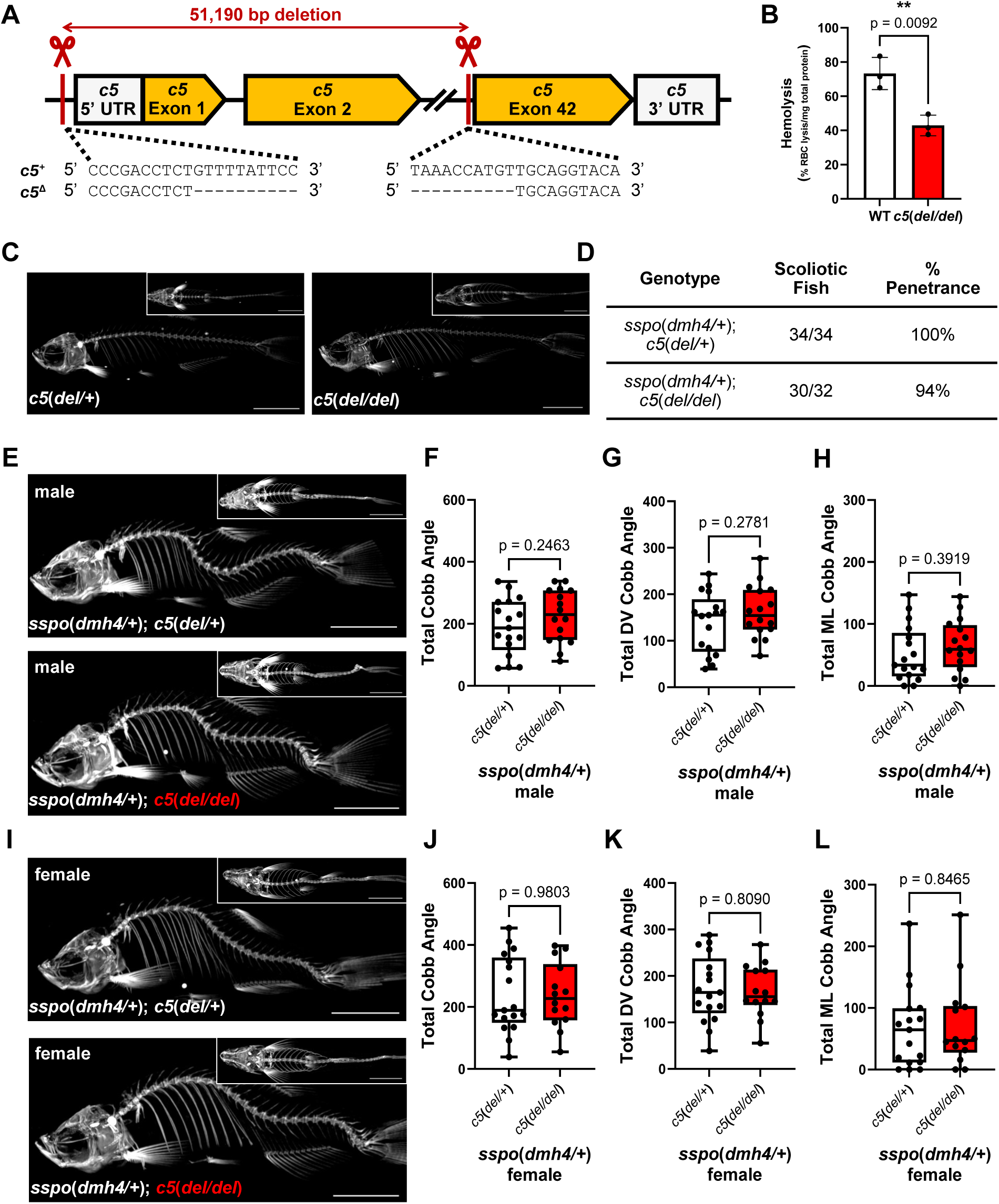
Loss of *c5* does not affect spinal curve severity in *sspo*(*dmh4/+*) IS models. (A) A large deletion of the *c5* ORF was generated, eliminating the 5’UTR and transcription start site. (B) Complement hemolysis assay functionally validates loss of C5 activity. (C) Loss of *c5* does not cause axial phenotypes in a wildtype background. (D) Quantification of scoliosis penetrance in *sspo*(*dmh4/+*); *c5*(*del/+*) and *sspo*(*dmh4/+*); *c5*(*del/del*) siblings revealed no significant difference between cohorts (Two-sided Fisher’s exact test, p=0.2312; if separated by sex: females, p=0.4687; males, p>0.9999). To quantify the effect of *c5* deletion on *sspo*(*dmh4/+*) scoliosis severity in male (E) and female (I) fish, uCT imaging was performed, followed by Cobb angle analysis (F-H, J-L). The scoliosis phenotype was not penetrant in two fish (genotype: *sspo*(*dmh4/+*); *c5*(*del/del*), 1 male and 1 female). These data points have been eliminated from the analysis of scoliosis severity. Scale bars = 5mm.

Furthermore, Cobb angle analysis revealed no difference in scoliosis severity between these groups in either plane of rotation (Fig. 5 E-L). Hence, we concluded that loss of *c5*, and therefore loss of MAC activity, is not necessary for scoliosis incidence or scoliosis progression in the *sspo(dmh4/+)* IS model.

### Deletion of *c5* strongly increases scoliosis severity in male *ptk7a*(*hsc9/hsc9*) fish

Shifting our focus to the nuances of zebrafish IS models, *ptk7a(hsc9/hsc9)* mutants were a unique candidate for our investigation of the role of *c5* in IS pathogenesis.

Firstly, this IS model presents with the strongest and broadest complement transcriptional signature that we are aware of, notably displaying upregulation of every terminal pathway gene *c5-c9* (Fig. S5). Also, this model uniquely experiences hydrocephalus which we expect triggers tissue regeneration pathways (Chen et al., 2024; Karimy et al., 2020), potentially implicating the regenerative functions of *c5* (Peterson and Anderson, 2014). In order to specifically study the endogenous role of complement in *ptk7a*(*hsc9/hsc9*) spine morphogenesis, we utilized the only available homozygous viable complement mutation: the *c5* deletion.

Scoliosis was fully penetrant in all genotypic groups within this clutch. Cobb angle analysis between *ptk7a(hsc9/hsc9)*; *c5(del/del)* fish and *ptk7a(hsc9/hsc9)*; *c5(del/+)* control siblings revealed no difference in scoliosis severity in either plane of rotation in females (Fig. 6E-H). Yet, in male *ptk7a(hsc9/hsc9)* mutants, we observed that a complete loss of *c5* lead to strikingly more severe spine curvatures in both planes of rotation (Fig. 6A-D).

**Figure 6:**
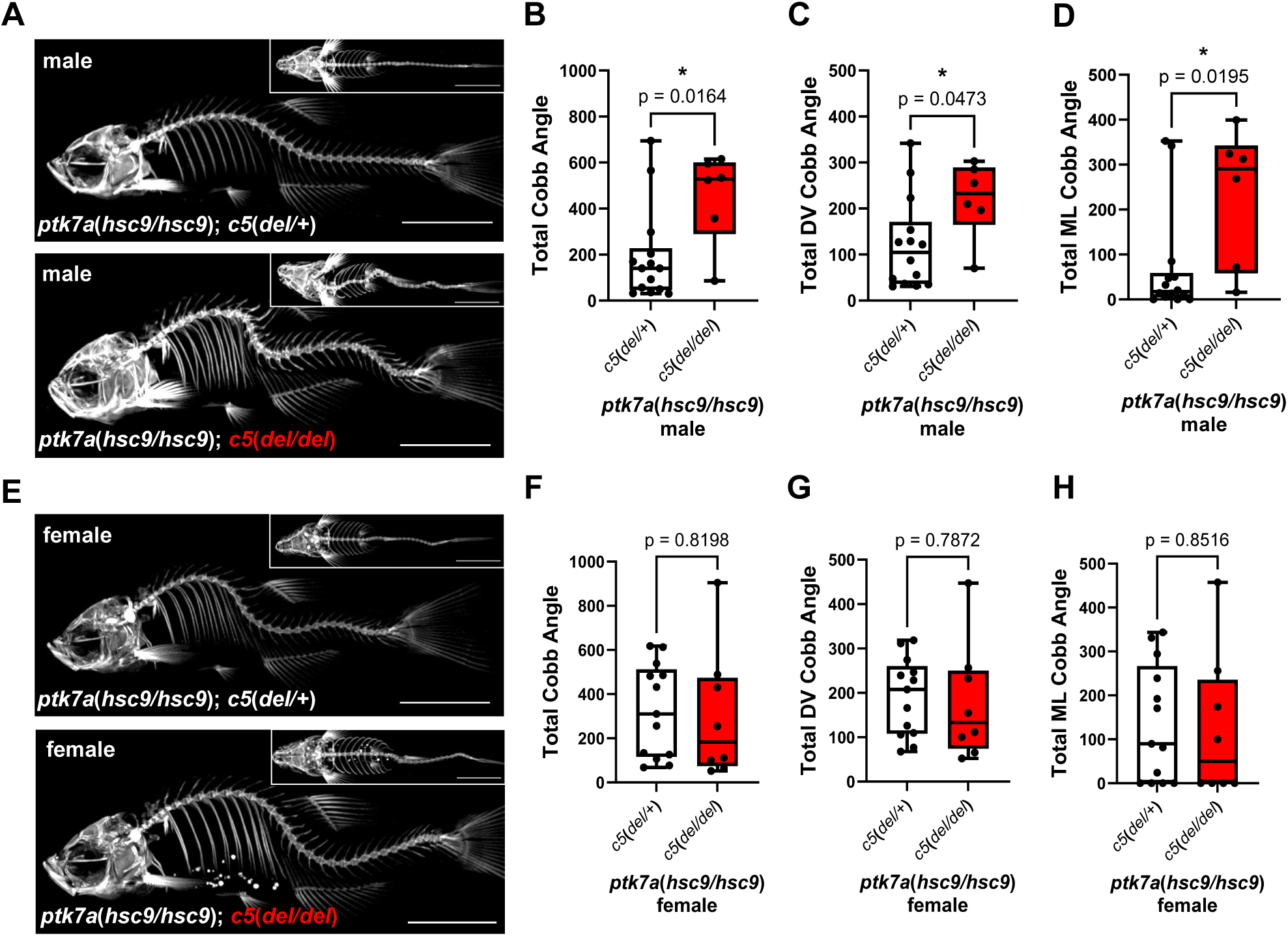
Loss of *c5* in the *ptk7a(hsc9/hsc9)* IS model leads to a male-specific increase in spinal curve severity. To quantify the effect of *c5* deletion on *ptk7a(hsc9/hsc9)* scoliosis severity, uCT imaging was performed (A, E), followed by Cobb angle analysis (B-D, F-H). The scoliosis phenotype was fully penetrant. Scale bars = 5mm.

Notably, we observed more severe spinal curvatures in *ptk7a(hsc9/hsc9)*; *c5(del/+)* female controls than in male counterparts, which is in line with previous documentation of sex bias in this model (Hayes et al., 2014). However, the complete loss of *c5* eliminated this sex bias (Fig. S6).

## DISCUSSION

In this study, we investigated the contribution of complement system activity to the etiology of idiopathic scoliosis using newly generated zebrafish mutants and stable transgenic lines targeting key complement components, C3 and C5. These tools enabled modulation of complement activity ubiquitously and within the central nervous system across established IS models. We demonstrate that pan-neuronal overexpression of *c3* by *Tg(elavl3::c3*) is not sufficient to induce scoliosis but exacerbates spinal curvature in female *sspo(dmh4/+)* mutants. In contrast, although membrane attack complex (MAC) activity is not required for scoliosis onset in either *sspo(dmh4/+)* or *ptk7a(hsc9/hsc9)* models, loss of *c5* markedly worsens spinal curve severity in male *ptk7a(hsc9/hsc9)* mutants. Together, these findings support sex-specific roles for complement activity in modulating scoliosis progression rather than disease initiation. Additionally, the stable genetic tools developed here provide a foundation for future studies of complement-mediated processes in embryonic and adult zebrafish.

### Implication of Complement Component 3 activity in IS severity

Although prior work reported no sex-associated difference in scoliosis severity in the *sspo(dmh4/+)* IS model (Rose et al., 2020), we observed that neuronal *c3* overexpression led to a modest but statistically significant increase in curvature severity specifically in female mutants. These findings suggest that scoliosis progression in the *sspo(dmh4/+)* model is sensitive to C3 levels in a sex-dependent manner.

We propose a threshold hypothesis, where it takes a smaller increase in inflammation to exacerbate scoliosis in females than in males. This concept aligns with human GWAS data suggesting that adolescent males may require a greater burden of genetic risk factors to develop IS, consistent with a lower susceptibility threshold in females (Kruse et al., 2012). Sex-associated differences in innate complement protein expression levels and activation levels have been previously documented in humans (Gaya da Costa et al., 2018; Kamitaki et al., 2020), although such differences remain uncharacterized in teleosts. Baseline sex-specific differences in C3 abundance and/or activation states could potentially underlie our findings and warrant direct investigation. Future studies should evaluate sex-segregated complement gene expression profiles in zebrafish IS models. Even in disease models lacking overt sex bias, our results underscore that genetic or inflammatory modifiers may differentially affect disease progression between sexes.

Importantly, overexpression of the complement regulator *rca2.1* [*Tg(βactin2::rca2.1)*] did not suppress scoliosis *in sspo(dmh4/+) fish*. This finding argues against a dominant role for C3-mediated opsonization in driving disease progression. Instead, the exacerbation observed with neuronal *c3* overexpression may reflect increased anaphylatoxin signaling (e.g., C3a activity), although mechanistic validation is required. This interpretation is consistent with previous reports that endogenous *c3* expression is upregulated in *sspo(dmh4/+)* animals and proposed to be pathogenic (Rose et al., 2020).

It is important to consider that endogenous *c3* upregulation in the *sspo(dmh4/+)* model is modest (supplemental figure 2) and may represent a secondary consequence of disease. The absence of phenotypic rescue in *Tg(βactin2::rca2.1)* fish is compatible with this possibility. However, this does not contradict our finding that exogenous neuronal *c3* overexpression exacerbates scoliosis, as forced elevation of *c3* is expected to increase inflammation within the nervous system independent of endogenous *c3* gene regulation. Our ability to interrogate endogenous C3 function is currently limited by the presence of eight *c3* paralogues in zebrafish and the lethality associated with homozygous *rca2.1* loss. Within these constraints, we conclude that neuronal *c3* upregulation is sufficient to worsen scoliosis in female *sspo(dmh4/+)* mutants, revealing a sex-specific vulnerability in this model.

### Implication of Complement Component 5 in IS Severity

Loss of terminal pathway activity, via *c5* deletion, had no measurable impact on scoliosis in *sspo(dmh4/+)* mutants but significantly increased spinal curve severity in male *ptk7a(hsc9/hsc9)* fish. It is intriguing that *c5* loss had a detrimental effect, that it affected only one of the two IS models, and that it aggravated curvatures specifically in male fish. This result was unexpected, as our initial hypothesis predicted that complement activation would contribute pathologically to scoliosis progression. Instead, these findings suggest that C5 and terminal complement activity may play a protective or compensatory role in maintaining axial straightness in the *ptk7a* model.

The model-specific effect of *c5* deletion highlights pathophysiological differences between *sspo(dmh4/+)* and *ptk7a(hsc9/hsc9)* IS models. Notably, complement transcriptional activation is stronger and more widespread in the *ptk7a* model (supplemental figures 2 and 5), suggesting distinct regulatory feedback dynamics that may differentially sensitize each model to terminal pathway disruption (Mason et al., 1999). Another distinction is that hydrocephalus has been observed in the *ptk7a* model (Grimes et al., 2016). Hydrocephalus triggers acute inflammation which is linked to tissue repair (Chen et al., 2024; Karimy et al., 2020). Given that tissue regeneration functions of complement have been widely documented, we might further extend them to the context of hydrocephalus-induced tissue damage (Peterson and Anderson, 2014). Therefore, alterations in complement activity could negatively affect tissue repair in the *ptk7a* model, resulting in more severe scoliosis.

The sex-specific effect of *c5* loss further supports our proposed complement threshold model. In *ptk7a* mutants, females exhibit more severe baseline scoliosis than males. It is therefore plausible that female mutants approach a maximal viable curvature severity, limiting the detectable impact of *c5* deletion. In contrast, male mutants may remain below this ceiling, allowing *c5* loss to drive progression toward a more severe phenotype comparable to that observed in females. These findings implicate C5-derived effectors (C5a and/or C5b) in maintaining axial integrity in a sex-dependent manner, though further mechanistic studies are required.

### Beyond Complement: Inflammation and Early Disease Mechanisms

Growing evidence from clinical and experimental studies implicates inflammation as a hallmark of IS (Bertelè et al., 2024; Djebar et al., 2024; Makino et al., 2019; Rose et al., 2020; Shen et al., 2019; Van Gennip et al., 2018; Wang et al., 2021). However, most clinical investigations focus on systemic inflammatory markers in blood, leaving the tissue-specific sources, timing, and mechanistic consequences of inflammation largely unresolved. Access to relevant spinal tissues in humans is typically restricted to surgical biopsies from severe cases, limiting temporal and etiological insight. Moreover, clinical diagnosis occurs only after curvature onset, as predictive biomarkers are lacking.

Animal models are therefore essential for dissecting early pathogenic events and distinguishing primary drivers from secondary or compensatory processes. The present work contributes to this effort by functionally interrogating complement activity within defined genetic models of IS.

Inflammation does not operate in isolation. Oxidative stress and other cellular stress pathways are tightly interconnected with immune signaling. Recent work demonstrated that oxidative stress–induced extracellular matrix defects precede scoliosis onset in the *sspo(dmh4)* model (Pumputis et al., 2025). Because oxidative stress can both trigger and result from complement activation (Hart et al., 2004; West and Kemper, 2023), reciprocal amplification loops may occur. Temporal dissection of redox imbalance and complement dysregulation will be critical to distinguishing early initiating events from downstream progression mechanisms.

### Conclusions and Future Directions

Our study identifies complement components C3 and C5 as modulators of scoliosis progression in zebrafish IS models in a sex-biased manner. While mechanistic pathways remain to be elucidated, our findings emphasize the importance of incorporating sex as a biological variable in preclinical IS research.

We also report the development of stable genetic tools for manipulating complement activity in zebrafish, enabling future studies aimed at defining conserved and species-specific aspects of complement biology. Given that complement signaling contributes to sex-biased susceptibility in multiple human diseases (Kamitaki et al., 2020), it is plausible that complement activity may also influence the greater severity and prevalence of IS observed in female patients. If translational parallels are confirmed, complement-targeted therapeutics (Mastellos et al., 2019) could represent a novel avenue for modifying disease progression. However, careful mechanistic dissection will be required before such strategies can be rationally pursued.

## Acknowledgements

We gratefully acknowledge the SickKids’ Zebrafish Facility technicians for excellent zebrafish care, as well as Jenica van Gennip and Matthew Mabey for technical assistance. This work was supported by the Canadian Institutes of Health Research Foundation grant FDN-167285 and the Canada Research Chair Program (B.C.).

## Materials and Methods

### Animal care

Established *Danio rerio* (zebrafish) husbandry protocols were adhered to and performed in accordance with Canadian Council on Animal Care (CCAC) guidelines. Wildtype zebrafish from TU strains were used. The *ptk7 hsc9* (Hayes et al., 2013) and *sspo dmh4* (Henke et al., 2017) alleles used in this study have been previously described. Embryos from natural matings were grown at 28°C. When required, experimental animals were euthanized with tricaine (500 mg/L; MS-222/MESAB), followed by submersion of anesthetized fish in ice water for several minutes.

### Molecular cloning

#### Cloning *c3b.2*

A cDNA library was prepared from adult *ptk7*(*hsc9/*hsc9) gut tissue RNA using SuperScript IV reverse transcriptase and oligo dT (Invitrogen). The full-length open reading frame (ORF) of *c3b.2* was cloned using Phusion polymerase (NEB) following manufacturer’s instructions (primers F: 5’-ATGCTTCTTCAGCTGCTGTTA-3’; R: 5’-TCAAGCCTGACAGCCGTCTTC-3’). The amplicon size was verified by gel electrophoresis, isolated by gel extraction (NEB), and a secondary PCR was run to add attB1, Kozac, and attB2 sequences to the amplicon. Once isolated, the amplicon was recombined into a pDONR221 vector using BP clonase (Gateway, Invitrogen).

#### Cloning *rca2.1*

A cDNA library was prepared from 1dpf embryo RNA as described above. A full length ORF of *rca2.1* was cloned (primers, F: 5’-ATGCCAAGAAATGTGCTTTCTCT-3’; R: 5’-TTAAAGCGATCCCTCTTCATTTG-3’) and a pME-*rca2.1* vector was generated.

### Transgenesis

Transgenes were assembled using standard Tol2 kit Gateway-compatible vectors via LR reactions (Gateway, Invitrogen). To generate Tg(*elavl3::c3b.*2) zebrafish, p5E-elavl3 (Addgene #75025), pME-c3b.2, and p3E-polyA were recombined into pDEST Tol2 HR2 transgenesis vector. To generate Tg(*Bactin2::rca2.*1) zebrafish, p5E-Bactin2, pME-rca2.1, and p3E-pA were recombined into pDEST Tol2 HR2 transgenesis vector.

Embryos were injected at the one-cell stage with 25 pg of assembled transgene and 25 pg of Tol2 mRNA. Embryos were sorted at 48 hpf for reporter expression (mCherry+ hatching glands) and were subsequently grown to adulthood. Individuals were bred to TU wildtype zebrafish to generate stable F1 lines. Subsequent F1 lines harboring the desired transgene were further outcrossed to TU wildtype zebrafish for line maintenance. Multiple independent lines of each transgene were generated, and expression levels were compared via qRT-PCR.

### CRISPR-Cas9 Mutagenesis

Large deletions across the ORF and smaller insertion/deletion mutations near the beginning of the gene were generated by co-injection of sgRNAs targeting both 5’ and 3’ ends of the genomic locus. sgRNAs were synthesized using sgRNA Synthesis Kit, S. pyogenes (EnGen) and purified using Monarch RNA Purification Kit (NEB). The 5uL injection mix was composed of 1uL Cas9 nuclease protein (EnGen, 20uM), 1uL KCl (2uM), and 150ng of each sgRNA (with up to 2 sgRNAs targeting either end of the gene). The mix was incubated at 37°C for 10 minutes, and then 1nL of mix was microinjected into one-cell stage embryos. At 2dpf, DNA was extracted from a subset of injected and non-injected control embryos, PCR amplification was performed around each site of cleavage, and the efficiency of CRISPR-editing was validated by Sanger sequencing using TIDE analysis (Brinkman and van Steensel, 2019).

To establish stable lines of fish with ORF-wide deletions, F0 injected fish were raised to adulthood, sperm extraction was performed on male adults, sperm DNA was extracted (Meeker et al., 2007), and PCR amplification was performed on each sample with the forward primer upstream of the targeted 5’ cut site and the reverse primer downstream of the 3’ cut site. In this screening method, the PCR extension time was limited to selectively amplify short amplicons resulting from the desired large deletion. Once founders were identified, they were outcrossed to Tu fish and the F1 generation was raised to adulthood and genotyped for desired mutations.

### *c5* deletion

sgRNA sequences: 5’-ATGAAGCAAACAGTCTATAA-3’ and 5’-GGGAACTTGTACCTGCAACA-3’.

Primers used for TIDE analysis amplifying 5’ end of gene F: 5’-ATGACAAGAACATTGCGCAATCA-3’, R: 5’-TTGACTGTAGTGACCGGGATTTT-3’; amplifying 3’ end of gene F: 5’-AAAGCAAAATCCATTGAGGAAA-3’, R: 5’-AGAATACTAGTTGCTGGCACCC-3’.

### *rca2.1* deletions

sgRNA sequences: 5’-GCGAGAGAAAGCACATTTCT-3’, 5’-TGGCATCTTATTTCGATGAA-3’, and 5’-CCTGACATGCAGTGAACTTG-3’.

Primers used for TIDE analysis amplifying 5’ end of gene F: 5’-GTCTTGATTGTTATGCTAGCCG-3’, R: 5’-CACAATTCATTCATGTTGTCCC-3’; amplifying 3’ end of gene F: 5’-TGCATAACAGACGTGCGAAT-3’, R: 5’-TTTTGGAGGCCGGTATATTG-3’.

### *rca2.1* and *rca2.2* co-deletion

sgRNA sequences: 5’-GAGATTATTTAAAACTCCCA-3’, 5’-TCGTATTTATATGAGGACAA-3’, and 5’-CCTGACATGCAGTGAACTTG-3’

Primers used for TIDE analysis amplifying 5’ end of gene F: 5’-CCCATCCTGAAAGGAAAATATC-3’, and R: 5’-AAAAAGGCTGAATAACAGCTCA-3’; amplifying 3’ end of gene: same as for *rca2.1* deletion.

### Genotyping

#### *c5* mutant

Triple primer PCR using F: 5’-ATGACAAGAACATTGCGCAATCA-3’, R: 5’-TTGACTGTAGTGACCGGGATTTT-3’, and R: 5’-AGAATACTAGTTGCTGGCACCC-3’.

Annealing temperature of 60°C and 55 second extension time (1 min/kb).

#### *rca2.1* mutants

*rca2.1 del* – Triple primer PCR using F: 5’-ATGCTAGCCGCTAACAGAAAA-3’, R: 5’-GATAGATAGCTACTACTCACCT-3’, R: 5’-TTTTGGAGGCCGGTATATTG-3’. Annealing temperature of 60°C and 30 second extension time (1 min/kb).

*rca2.1* 3bpΔ and 4bp Δ - High resolution melting analysis (Light Cycler 96 Instrument, Roche) using F: 5’-ATTAATTGCAGAAATCAAAAACG-3’ and R: 5’-CAAAAGACAGGAGAACAAAGCA-3’.

#### rca2.1-rca2.2 mutant

Triple primer PCR using F: 5’-CCCATCCTGAAAGGAAAATATC-3’, R: 5’-AAAAAGGCTGAATAACAGCTCA-3’, and R 5’-TTTTGGAGGCCGGTATATTG-3’. Annealing temperature of 60°C and 1:30 minute extension time (1 min/kb).

### Micro-CT imaging, image processing, and Cobb angle analysis

Zebrafish were euthanized at 4mpf and fixed in 10% neutral-buffered formalin (Sigma-Aldrich) for at least 24 hours. Fish were mounted in tubes using 1% agarose. Scanning was performed with a SkyScan 1275 microCT (Bruker, Kontich, Belgium) using 50 kV and 85 μA, sample rotation of 180°, image rotation steps of 0.2°, frame averaging of 3, camera binning of 1×1, and using a pixel size of 21-45 um.

Projection images were reconstructed into cross-sections using SkyScan’s NRecon v.1.7.4.6 software (Bruker, Kontich, Belgium) in a range of attenuation coefficients 0–0.2, with a beam-hardening correction of 40%. Images were manipulated using CTvox software (Bruker microCT) to obtain images of each fish from lateral and dorsal views for subsequent Cobb angle analyses.

Cobb angles were measured using SCODIAC software (Pavel Cerny and Lukasz Stolinski) as demonstrated in supplemental figure 1. Lines were drawn at the distal edge of the top and bottom most displaced vertebrae for each curve. The Cobb angle was then measured as the angle of intersection between lines drawn perpendicular to the original 2 lines. Cobb angle measurements from lateral and dorsal images were summed to achieve a total Cobb angle value. Prism software (GraphPad) was used to perform unpaired two-tailed t-test analysis of scoliosis severity.

### Quantitative reverse transcription polymerase chain reaction

Total RNA was extracted from 10 larva (5dpf) using TRIzol reagent (Invitrogen). qRT-PCR for validation of transgene overexpression was performed using 10 ng of total larval RNA in a one-step qRT-PCR (Luna Universal One-Step RT-qPCR Kit, NEB) and performed on a Roche LightCycler 96 system. The following primers were designed and used: GAPDH F: 5’-GGGCTGCCAAGGCTGTAGGC-3’ R: 5’-TGGGGGTGGGGACACGGAAG-3’; *rca2.1* F: 5’-GGTGGAATTTTCGGTAGTGG-3’ R: 5’-CCCTCTTCATTTGTAGGCACC-3’; *c3* F: 5’-ATGCTTCTTCAGCTGCTGTTA-3’ R: 5’-TGTCCCTCCACCAGGATGTTC-3’. All graphs are representative of three independent experiments with three technical replicates.

For qRT-PCR analysis, Ct values were obtained for target genes and normalized to GAPDH. Fold change was calculated relative to wildtype expression according to the following equation: 2^−ΔΔCt^. Statistical significance was determined by unpaired two-tailed t-test using GraphPad Prism.

### Hemolysis assay

Complement activity (MAC-induced lysis) was measured as the level of red blood cell (RBC) lysis by zebrafish whole body homogenate (WBH). The experimental protocol was based on previous published protocols (Oriol Sunyer and Tort, 1995; Wang et al., 2009). Rabbit RBCs were obtained from Innovative Research and GVBo buffer (without Ca and Mg) was obtained from Complement Technology Inc. WBH was isolated from WT and *c5*(*del/del*) larva (n=200 larva per sample, 6 dpf). BCA protein assay kits (Thermo Scientific, Pierce) were used to standardize protein levels across samples. The assay was run as 3 independent biological replicates with 3 technical replicates. Statistical significance was determined by unpaired two-tailed t-test using GraphPad Prism.

---

**Supplemental figure 1:**
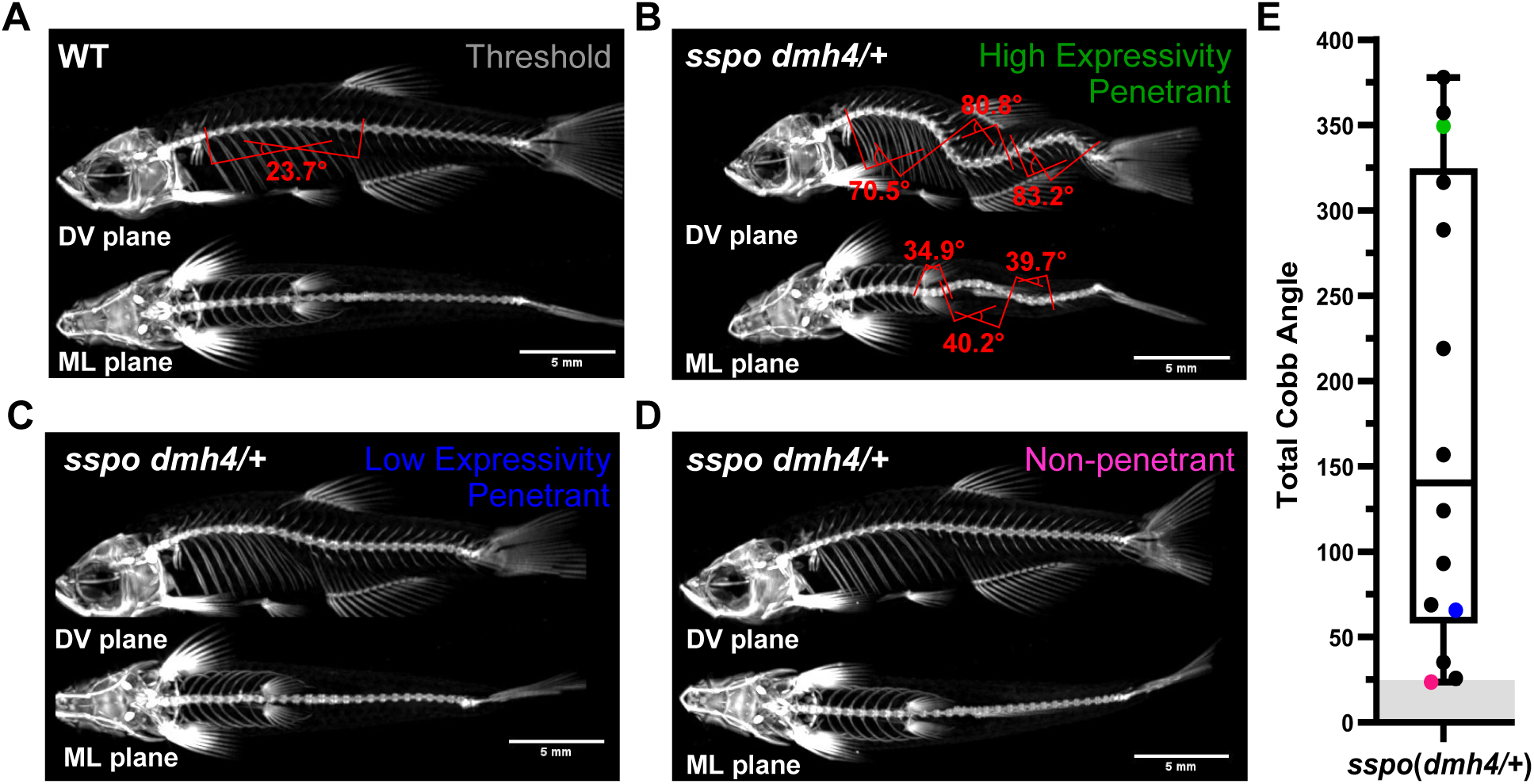
Sex-specific Cobb value thresholds enable objective classification of penetrance in the *sspo(dmh4/+)* IS model. (A) uCT image of an adult male wildtype zebrafish that exhibits a natural 24° spinal curvature, which represents the upper threshold before diagnosis of scoliosis. (B-D) uCT images of adult male *sspo(dmh4/+)* siblings that exhibit varying degrees of scoliosis penetrance and expressivity. (E) Distribution of Cobb angle measurements in male *sspo(dmh4/+)* zebrafish, with threshold of scoliosis diagnosis indicated (grey box).

**Supplemental figure 2:**
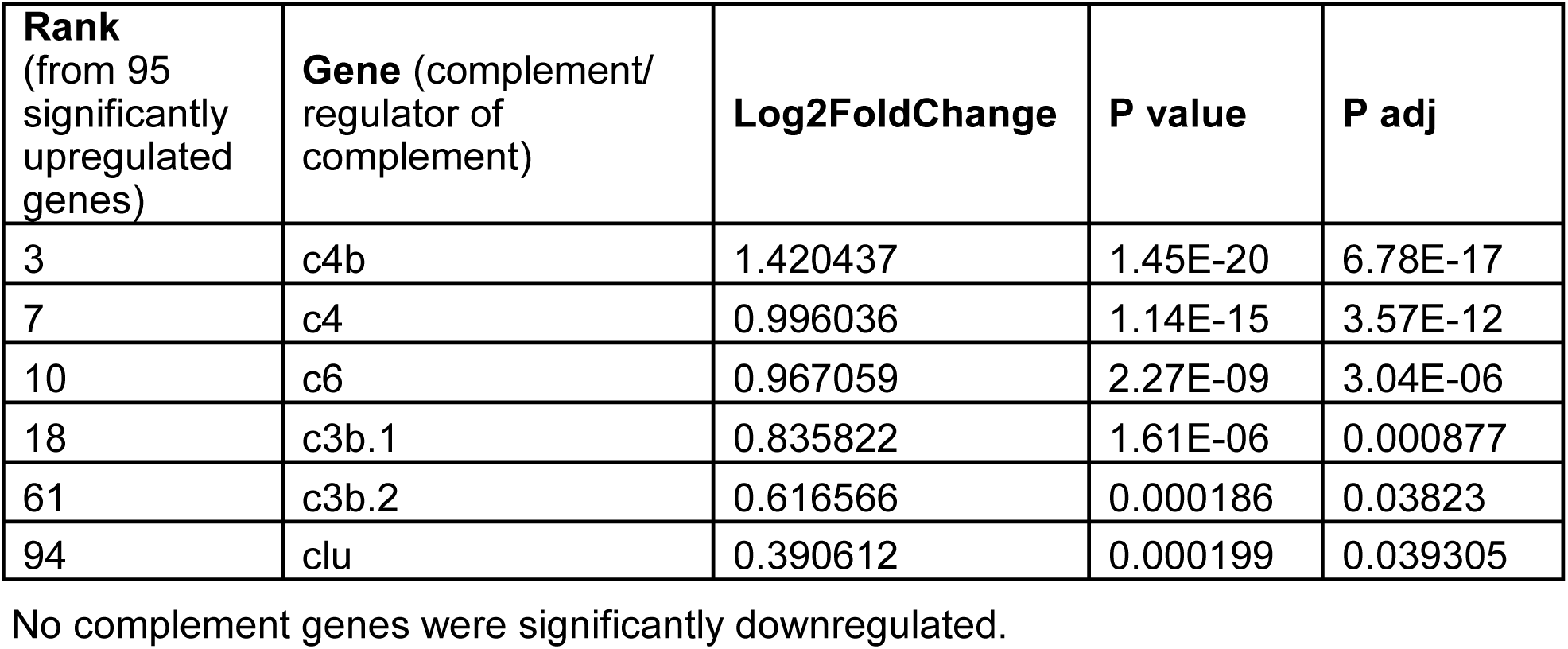
Genes associated with the complement pathway are significantly upregulated in *sspo(dmh4/+)* IS model. Data obtained from RNA-sequencing of whole brains of *sspo(dmh4/+)* fish with severe scoliosis vs. wildtype siblings, brains were dissected at 3wpf, post scoliosis onset (from Rose et al., 2020).

**Supplemental figure 3:**
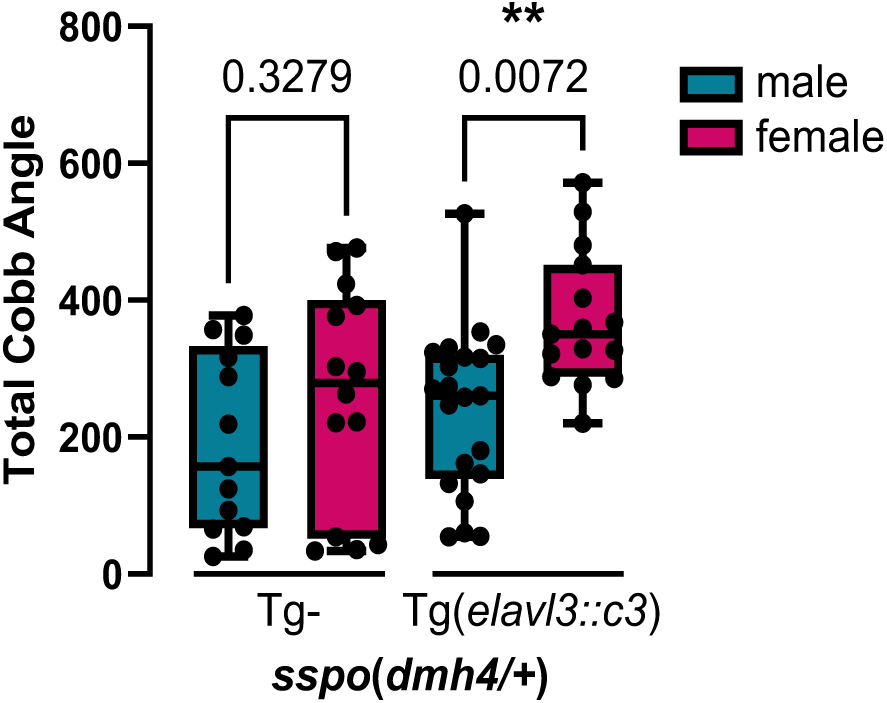
Female *sspo(dmh4*/+); Tg(*elavl3::c3*) fish exhibit a more severe total Cobb angle value than male siblings. Statistical significance measured by one-way ANOVA test.

**Supplemental figure 4:**
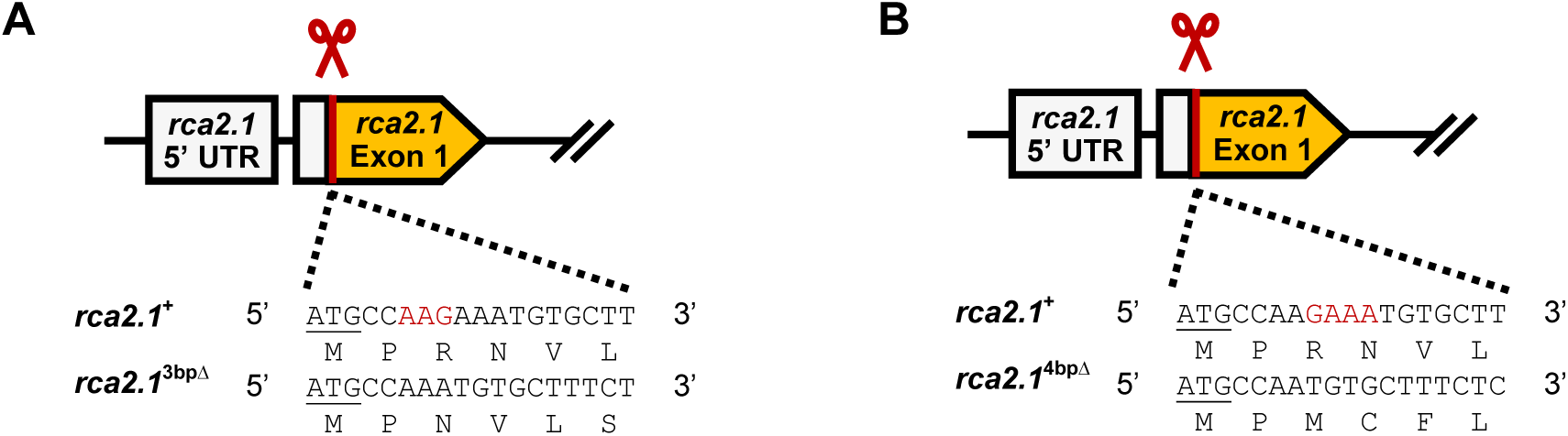
Stable in-frame and out-of-frame *rca2.1* mutant alleles were generated. Both mutations are homozygous lethal. (A) A 3bp insertion is located within the signal peptide sequence (Ensembl) and predicted to disrupt membrane localization. (B) If a stable transcript results from this frame-shift mutation, the predicted protein product would be: MPMCFLSLLLCSPVF*.

**Supplemental figure 5:**
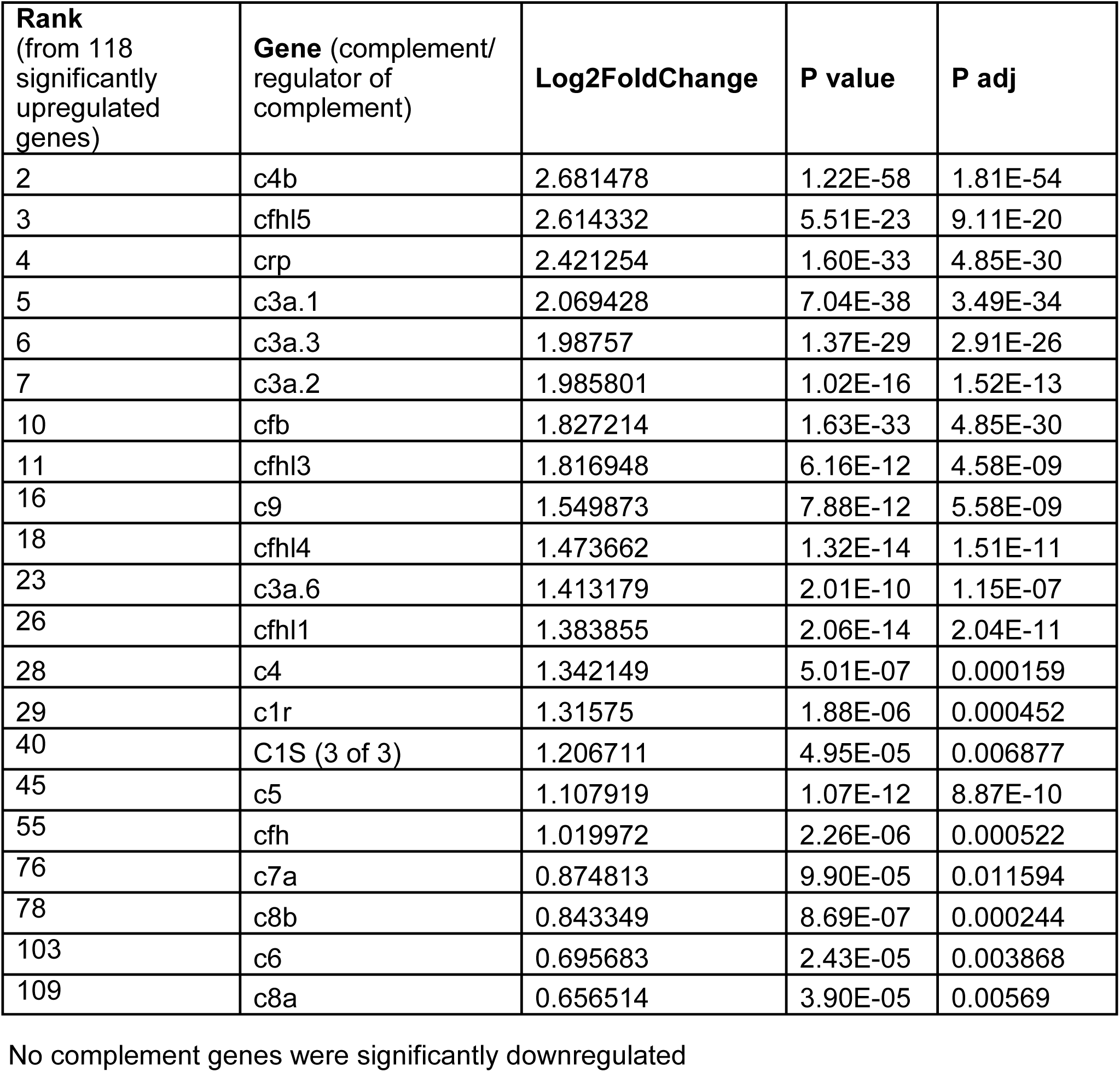
Genes associated with the complement pathway are significantly differentially expressed in *ptk7a(hsc9/hsc9)* IS models. Data obtained from RNA-sequencing of whole trunks and tails of *ptk7a(hsc9/hsc9)* fish with severe scoliosis vs. *ptk7a(hsc9/+)* siblings (post spine curvature onset) (from Van Gennip et al., 2018).

**Supplemental figure 6:**
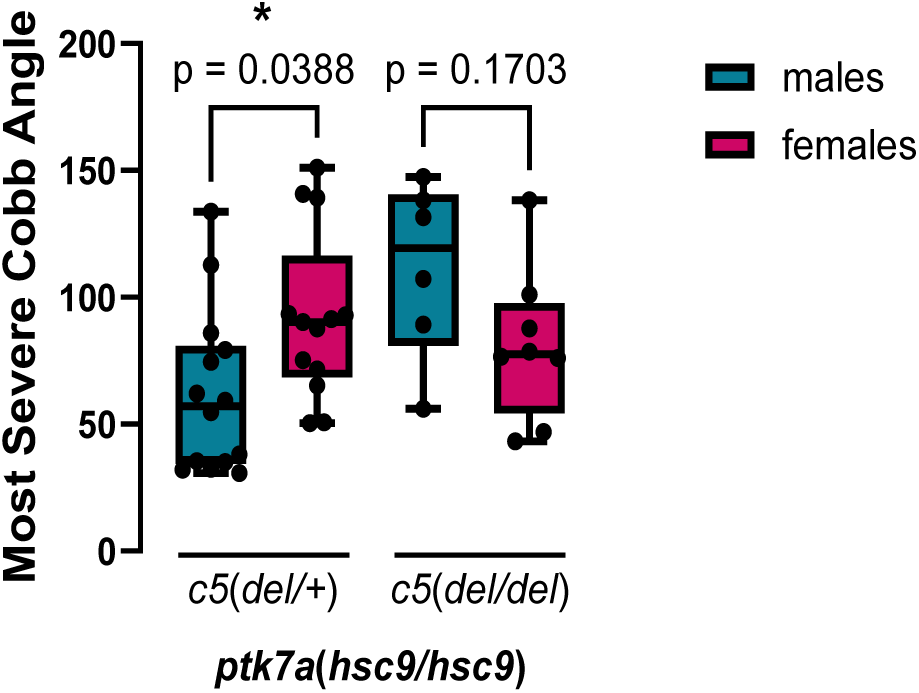
Homozygous loss of *c5* eliminates existing sex-bias of most severe spinal curvatures in *ptk7a*(*hsc9/hsc9*) IS models. Statistical significance measured by one-way ANOVA test.

